# DNA damage hypersensitivity in chicken primordial germ cells limits CRISPR-Cas9 editing and supports an shRNA-based screening approach

**DOI:** 10.1101/2025.09.17.676926

**Authors:** Chen Kwoon Zhang, Xin Zhiguo Li

**Author notes:** Correspondence (X.Z.L.).

## Abstract

Chickens (*Gallus gallus*) are uniquely suited for germline studies because their primordial germ cells (PGCs) can be propagated long-term in vitro and used for germline transmission. To develop a loss-of-function screening platform in chicken PGCs, we compared three perturbation methods: CRISPR/Cas9 knockout, CRISPR interference (CRISPRi), and shRNA-mediated knockdown. We found that CRISPR/Cas9 editing causes severe toxicity in PGCs, with DNA damage hypersensitivity over 20-fold greater than in somatic cells, and with distinct DNA damage checkpoint responses between male (ZZ) and female (ZW) lines. CRISPRi using dCas9-KRAB was ineffective in chicken—likely because of species-specific epigenetic constraints—whereas shRNA knockdown produced robust, nontoxic gene silencing. These results identify DNA damage hypersensitivity as a major barrier to nuclease-based editing in the germline and establish RNAi as a feasible platform for genome-wide functional screening in chicken PGCs.

## Introduction

Chickens (*Gallus gallus*) have long been more than agricultural mainstays – they are a key model organism in biological research. The chick embryo system has a distinguished history in developmental biology and has contributed fundamental insights in immunology, genetics, virology, and cancer research(Stern, 2005). Several features make the chicken a particularly advantageous model: it is an amniote with accessible embryonic development, a fully sequenced genome, and it is amenable to diverse experimental manipulations(Stern, 2005). These attributes, combined with emerging genetic tools, have solidified the chicken’s status as an ideal model for studies ranging from evolutionary biology to biomedical applications. Indeed, breakthroughs enabling transgenic and genome-edited chickens have transformed the chicken from primarily a correlative model to a direct causal one in experimental genetics(Salter et al., 1987; Schusser et al., 2013; Park et al., 2014). This rich legacy underlines that beyond its commercial importance, the chicken is an invaluable system for uncovering basic biological principles.

A major technical advantage of the chicken model is the ability to culture primordial germ cells (PGCs) long-term in vitro, a feat unique among vertebrates(Karagenç et al., 1996). PGCs are the embryonic precursors of sperm and eggs, and being able to propagate these cells indefinitely in culture has transformative implications. Stable chicken PGC lines retain their germline identity and competency over extended periods(Karagenç et al., 1996), enabling researchers to perform genetic modifications and then produce live transgenic offspring from those edited cells(Park and Han, 2012; Oishi et al., 2016; Dehdilani et al., 2023). Indeed, establishing long-term PGC cultures has been crucial for creating transgenic lines, including bioreactor chickens that deposit recombinant proteins in egg whites, and for preserving valuable poultry genetics. Notably, chicken is currently the *only* vertebrate species in which PGCs can be stably expanded in this manner(Karagenç et al., 1996). In contrast, mammalian PGCs cannot be maintained in an undifferentiated state long-term – they either spontaneously differentiate or die, necessitating alternative approaches. For example, mouse or human PGCs can be studied by *deriving* PGC-like cells (PGCLCs) from pluripotent stem cells in vitro, but not by continuously culturing the actual PGCs(Hwang et al., 2023). This makes the avian system uniquely powerful for germline genetic engineering and fundamental research. As a practical illustration, CRISPR/Cas9 editing of cultured chicken PGCs has been used to generate gene-edited chickens resistant to viral diseases like avian influenza(Idoko-Akoh et al., 2023). Such applications highlight how the chicken PGC platform enables fundamental research utility with agricultural and biomedical innovation.

Despite these advantages, several fundamental questions about germ cell biology remain that chickens are well-poised to address. PGCs must solve unique biological problems as the cells that carry genetic and epigenetic information between generations. For instance, how germ cells achieve dosage compensation of sex chromosomes and undergo sex- specific development is not fully understood. Birds have a ZZ/ZW sex chromosome system and incomplete dosage compensation, raising intriguing contrasts with the XX/XY system of mammals(Melamed and Arnold, 2007; Zimmer et al., 2016). Another critical aspect is the piRNA pathway and transposon regulation: germ cells across animal phyla rely on Piwi-interacting RNAs to silence transposable elements and protect genome integrity, yet many specifics of this pathway and its conservation are unresolved. Notably, disrupting key Piwi genes in mice leads to transposon de-repression and catastrophic germ cell loss due to apoptosis(Sun et al., 2022), underscoring the importance of this pathway. Many features of germ cell biology – from epigenetic reprogramming to meiosis initiation – are evolutionarily conserved, but others are species-specific or sexually dimorphic, necessitating a comparative approach. The availability of a long-term chicken PGC culture system now provides a unique opportunity to tackle these questions through unbiased, genome-wide functional screens conducted directly in bona fide germ cells.

In mammalian models, where stable PGC lines are unavailable, researchers have turned to surrogate in vitro systems to probe germ cell development. A prominent strategy has been to differentiate embryonic stem cells into primordial germ cell-like cells (PGCLCs), which mimic early PGCs, and then apply large-scale genetic perturbation screens in these PGCLC systems(Hwang et al., 2023). This approach has proven fruitful: in one high-throughput CRISPR interference screen targeting epigenetic regulators, inhibition of over 50 genes (such as the histone deacetylase co-repressor NCOR2) was found to impair mouse PGCLC formation, demonstrating the power of systematic screens to reveal germline developmental regulators(Li et al., 2025). Similarly, a genome-wide CRISPR knockout screen in human PGCLCs uncovered novel factors (e.g. the AKT–mTOR pathway activator TCL1A) that are crucial for human PGC-like cell development(Hwang et al., 2023). These studies highlight the utility of functional genomics in deciphering germ cell biology. However, PGCLCs remain an imperfect proxy for genuine PGCs. In fact, PGCLCs often fail to recapitulate later germline programs – for instance, mouse PGCLCs do not express key PIWI proteins and consequently lack piRNAs, unlike their in vivo PGC counterparts(Ramakrishna et al., 2022). This limitation means that certain germline phenomena (such as piRNA pathway activation, or the full sequence of epigenetic reprogramming events) cannot be faithfully studied in PGCLCs (Ramakrishna et al., 2022). Therefore, while PGCLC-based screens have been informative, there is a clear need to perform genome-wide screens in true PGCs to capture the complete biology. The chicken PGC system is ideally positioned to fill this gap, yet to date, such comprehensive screens in PGCs have lagged behind those in mammalian models. A major hurdle has been technical: establishing an efficient, high-throughput gene perturbation method in PGCs that does not compromise their survival or germline potential.

Here, we set out to develop a robust genome-wide screening approach for cultured chicken PGCs and to determine the most effective loss-of-function perturbation strategy for these sensitive cells. We evaluated three platforms side by side – CRISPR/Cas9 knockout, CRISPR interference (CRISPRi) using a dCas9-KRAB repressor, and short-hairpin RNA (shRNA) knockdown – by testing each method’s ability to induce loss-of-function phenotypes in chicken PGCs. Strikingly, conventional CRISPR/Cas9 genome editing proved highly deleterious to PGCs: Cas9-induced double-strand DNA breaks triggered extensive cell death, with PGCs being roughly tenfold more sensitive than somatic cells, and we observed a sex-specific bias in this toxicity. This hypersensitivity mirrors the p53- dependent cell-death response reported in human pluripotent stem cells after CRISPR editing(Ihry et al., 2018), and likely represents a protective mechanism guarding germline genomic integrity. Meanwhile, CRISPRi delivered inconsistent gene repression in PGCs and required sustained high dCas9-KRAB expression, limiting its utility. In contrast, shRNA-mediated knockdown emerged as a viable alternative, enabling effective high-throughput gene knockdown in PGCs without invoking acute DNA-damage toxicity. These findings reveal a pronounced DNA damage response in chicken PGCs and establish an shRNA-based framework for functional genomics in avian germ cells, paving the way for future applications in reproductive biology and biotechnology.

## Methods

### Chicken PGC Isolation and Culture

The eggs of Jinhua chicken used in this study were obtained from Yukang Agricultural Development Co., Ltd. Fertilized chicken eggs (*Gallus gallus*) were incubated at 37 °C (humidified, no rocking) until Hamburger–Hamilton stage 16 (60 h of incubation), when primordial germ cells (PGCs) enter the embryonic circulation. Embryos were accessed at the blunt egg pole using sterile forceps under a dissection microscope. The dorsal aorta was visualized and gently pierced with a glass capillary pipette to draw ∼1–2 µL of embryonic blood, which was immediately transferred into 200 µL of pre-warmed PGC culture medium in a 96-well plate. In some preparations, red blood cells were selectively lysed by mixing the blood with 10× volume of ACK lysis buffer (A1049201, Thermo) on ice for 15–30 min prior to seeding; remaining cells were pelleted (5 min, 200–300×g, 4 °C), washed with PBS, and resuspended in culture medium. The base culture medium was Avian KnockOut DMEM (a calcium-free modified DMEM) (PWL037, MeilunBio) supplemented with 1×B-27 (17504-044, Gibco), 1×non-essential amino acids (11140-050, Gibco), 1×GlutaMAX (35050061, Gibco), 1×nucleosides (ES-008-D, Millipore), 0.1 mM β-mercaptoethanol (M3148, Sigma), 1.34 mM sodium pyruvate (11360-039, Gibco), 0.15 mM CaCl₂ (499609, Sigma), 0.2% ovalbumin (A5503, Sigma), and 100ug/ml heparin sodium (HY-17567A, MCE). Prior to use, this basal medium was further supplemented with 100 µg/mL penicillin and streptomycin (15140122, Life), a mycozap (VZA-2022, Lonza), 4 ng/mL basic fibroblast growth factor (bFGF2, C046, novoprotein), 25 ng/mL Activin A (C687, novoprotein), 25 ng/mL insulin-like growth factor 1 (IGF-1,C032, novoprotein), 10 µg/mL chicken ovotransferrin (C7786, Sigma), and 0.2% filtered chicken serum (C2550-0100, VivaCell). This serum-free, low-calcium medium (sometimes termed “FAOT” medium for FGF2/Activin/Ovotransferrin) supports long-term expansion of chicken PGCs in semi-suspension culture (Whyte et al., 2015). Cultures were maintained at 37 °C in 5% CO₂, and half the medium was changed with the fresh medium every 2 days. PGCs remained mostly non-adherent, often forming small floating clumps; to prevent over- clumping, gentle pipetting was performed weekly after feeding. After about one week in culture, PGC colonies became visible. By 2–3 weeks, cultures typically contained several ×10^4^ PGCs, at which point they were expanded to larger wells or split to maintain a density below ∼3×10^5^ cells per 24-well (∼0.5 mL) to ensure logarithmic growth. Cultures with poor growth (< 5×10^4^ cells after 4 weeks) were considered non-established and discarded.

To cryopreserve PGC lines, cells in mid-log phase (approximately 1–2×10^5^ cells in 0.5 mL) were harvested and mixed 1:1 with filter-sterilized freezing solution (final 8% DMSO, 10% chicken serum in basal medium with 0.15 mM Ca²⁺). Aliquots were cooled in a controlled-rate freezing container at –80 °C overnight and transferred to liquid nitrogen. For revival, vials were thawed quickly at room temperature, and cells were immediately diluted dropwise into 4× volume of room-temperature basal medium. After centrifugation (5 min, 200×g), the cell pellet was resuspended in fresh PGC culture medium and returned to 37 °C. Typical post-thaw viability exceeded 90% for healthy cultures.

Each PGC culture was confirmed to consist of primordial germ cells by morphology and marker expression. Live PGCs were identified by their large size (> 12 µm) and distinctive phase-bright cytoplasm with refractive granules. For further confirmation, an aliquot of cells was examined by immunofluorescence staining for stage-specific embryonic antigen 1 (SSEA-1) and chicken VASA homolog (CVH, also called DDX4), which are specifically expressed by avian PGCs (See “Immunofluorescence” below for staining protocol). The sex of each PGC line was determined via PCR genotyping of the embryo’s genomic DNA: tissue from the same embryo was digested in alkaline lysis buffer, neutralized, and used as template for amplification of Z- and W-linked CHD1 gene fragments (580 bp Z allele; 434 bp W allele). No live animal procedures were performed in this study; all PGCs were obtained from early embryos (prior to hatching), and experiments were carried out entirely in vitro in compliance with institutional guidelines.

### Plasmid Transfection and Genome Editing in PGCs

For genetic manipulation of PGCs, we introduced plasmid DNA encoding various constructs (e.g. CRISPR/Cas9 and guide RNA expression vectors, CRISPR interference [CRISPRi] plasmids, or shRNA knockdown plasmids) into the cells using two complementary methods. First, a lipid-mediated transfection was tested on PGCs in suspension. Briefly, DNA (1 µg) was diluted in 50 µL Opti-MEM (31985070, Thermo Fisher) and combined with 3 µL of a transfection reagent FuGENE HD (E2311, Promega) prepared in 50 µL Opti-MEM. The DNA/reagent mixture was incubated ∼15 min at 37°C. PGCs were collected from culture (typically 4×10^4 cells per transfection), washed once with PBS, and resuspended in DNA/reagent mixture, then incubated ∼15 min at 37°C. After incubation, 100 µL cell/regent mixture were seeded in the 24 wells with 400 µL PGC medium. Transfected cells were cultured for 48 h, then assessed for transgene expression or subjected to antibiotic selection as needed. While convenient, we found that chemical transfection alone yielded relatively low efficiency in PGCs (typically < 20% EGFP⁺ cells), consistent with reports that standard lipofection is suboptimal in avian PGCs (Watanabe et al., 2023). The information on gRNA and shRNA is provided in Table S1. The RNA synthesis was performed by Shanghai Sangon Biotech, and the vector construction was carried out by Suzhou Zuoxing Biology.

To achieve higher transfection efficiency, we employed electroporation using a Lonza 4D- Nucleofector system. PGCs were concentrated by gentle centrifugation (5 min, 300×g) and resuspended in the Lonza P3 Primary Cell Nucleofector solution (20 µL per reaction, V4XP-3032, Lonza) with the plasmid DNA. For each electroporation, approximately 2×10^5^ PGCs and 1 µg of plasmid DNA were mixed in one well of a 16-well Nucleocuvette strip (Lonza). The cells were electroporated using Lonza® 4D-Nucleofector™ X Unit (AAF-1002X, Lonza) to program code CM-137. Immediately after the pulse, 100 µL of pre-warmed PGC medium was added to the cuvette, and the cells were gently transferred to a 96-well culture plate with an additional 100 µL of medium. This electroporation protocol is based on recent optimizations for feeder-free chicken PGCs and typically achieves high DNA delivery efficiency (≥ 60–70% of cells transfected) with good viability(Zhao et al., 2025). Following electroporation, PGCs were cultured for 48 h before downstream analyses or drug selection. For CRISPR/Cas9 gene editing, cells were electroporated with a mix of a Cas9/sgRNA plasmid (1-3 µg Cas9 at equivalent molar amounts [1:1.2 Cas9:sgRNA ratio]) ; co-transfection under these conditions has been shown to permit precise genomic integration in a substantial fraction of PGCs (Zhao et al., 2025).

To enrich for successfully transfected PGCs, we applied antibiotic selection corresponding to the resistance marker on the plasmid. After transfection (48 h), cells were pelleted and resuspended in PGC medium containing the appropriate antibiotic. For example, plasmids carrying a puromycin resistance gene were selected in puromycin (P8230, Solarbio) at 3 µg/mL for 12h. For blasticidin resistance, blasticidin S (B9300, Solarbio) was used at 40 µg/mL for 12h. These concentrations were determined by prior kill-curves on untransfected PGCs. After the selection period, surviving PGCs (presumed stably transfected) were expanded in normal PGC growth medium (without antibiotics) for use in assays.

### DNA Damage Treatment Assays

To examine DNA damage responses in PGCs, we treated cells with a DNA double-strand break (DSB)-inducing chemotherapeutic or ionizing radiation. For chemical induction of DSBs, PGC cultures were exposed to etoposide (33419-42-0, MCE) at a various concentrations of 0.01 µM, 0.1 µM, 1 µM, 10 µM, 100 µM, 1 mM for 24h and same concentration of etoposide for 48h for control cells. Etoposide (a topoisomerase II inhibitor) creates DNA breaks and activation of DNA damage signaling (Saleh et al., 2012), enabling us to assess repair responses. For physical induction of DNA damage, PGCs were subjected to X-ray irradiation using an X-ray generator (RS2000Pro-225 model, Radsource). Cells in suspension (in 96 well culture plates) were irradiated at a dose of 2,4,6,8 Gy (dose rate 1.5 Gy/min). Sham-irradiated controls were kept on room temperature for an equivalent duration. Following treatments, cells were either returned to the incubator for recovery (48h) before analysis. DNA damage was primarily quantified by immunofluorescent detection of phosphorylated H2AX (γ-H2AX) foci (see below). In addition, cellular viability after DNA damage was measured using the CCK-8 assay and cell cycle was evaluated by flow cytometric analysis of Annexin V/PI staining (see Cell Viability Assay below). All DNA damage experiments were performed in at least triplicate independent PGC cultures.

### Periodic Acid-Schiff (PAS) Staining

PGCs were first fixed with 70% ethanol for 10 minutes. PAS staining kit (G1360-5, Solarbio)was used, then performed as follows: 100 μl of periodic acid solution, equilibrated to room temperature, was added to each sample and incubated in a humidified chamber for 10 minutes in the dark. Cells were collected and washed 3 times in PBS (5 min each). Next, 100 μl of Schiff’s reagent was added, and the samples were incubated in a humidified chamber at 37°C in the dark for 30 minutes. After removing the reagent, the samples were washed in PBS for 5 minutes. This was followed by counterstaining with hematoxylin for 30 seconds and rinsing with PBS to remove excess dye. Images could be acquired directly by seeding in a 24-well plate.

### Immunofluorescence Staining and Imaging

For immunofluorescence analysis, approximately 5×10^4^ PGCs (which grow in suspension) were placed onto a glass slide (in a Pap pen-marked well) and allowed to settle and dry on a 37 °C slide warmer for ∼10 min, facilitating their attachment to the glass. Cells were then fixed in 4% paraformaldehyde/PBS for 10 min at room temperature and washed 3 times with PBS. Permeabilization was performed using PBS containing 0.1% Triton X-100 (and 0.01% Tween-20) for 20 min, followed by three PBS washes. Non-specific binding was blocked by incubating with 1% bovine serum albumin (BSA) in PBS for 1 h at room temperature. Primary antibodies were diluted in 3% BSA/PBS and applied to the cells overnight at 4 °C (or 2–3 h at room temperature). The following primary antibodies were used: anti-SSEA-1 (mouse monoclonal IgM, sc-21702, Santa Cruz Biotechnology, dilution 1:500), anti-VASA/DDX4 (rabbit polyclonal IgG, ab13840, Abcam, dilution 1:100), and anti-γ-H2AX (phospho-H2AX Ser139, 07-164, Millipore, dilution 1: 1000). After primary incubation, slides were washed 3× in PBS (5 min each). Appropriate secondary antibodies were then applied for 1–2 h at room temperature in the dark. These included Alexa Fluor® 488-conjugated secondary antibody against mouse IgM/IgG (for SSEA-1,a11029, Thermo Fisher, dilution 1:1000), Alexa Fluor 488 goat anti-rabbit IgG (H+L) (for γ- H2AX, a11008, Thermo Fisher, dilution 1:1000) and Alexa Fluor 594-conjugated anti-rabbit IgG (for VASA, a11012, Thermo Fisher, dilution 1:1000). After three final PBS washes, slides were mounted with fluorescence mounting medium containing DAPI (4′,6- diamidino-2-phenylindole, 1 µg/mL) to counterstain nuclei. Coverslips were sealed with nail polish and slides were stored at 4 °C in the dark until imaging.

Fluorescent images were acquired using a microscope (Evident (Olympus) SpinSR spinning disk confocal microscope) equipped with appropriate filter sets and a 60 × objective. Exposure settings were kept constant between control and experimental samples. PGC identity was confirmed by co-staining: PGCs typically showed strong SSEA-1 on the surface and cytoplasmic VASA signal, distinguishing them from any somatic cell contaminants. Negative control slides (no primary antibody) showed negligible fluorescence. All image processing and analysis were performed with Cellsens Dimension software under identical parameters for all groups.

### Cell Viability and Proliferation Assays

PGC proliferation rates and viability under various conditions were assessed using the Cell Counting Kit-8 (40203ES76, Yeasen Biotech) colorimetric assay. PGCs were plated in 96- well plates (flat-bottom) at ∼3×10^3 cells per well in 200 µL of PGC medium with treatments applied as described (such as DNA-damaging agents or other compounds, with appropriate vehicle controls). At the designated time points, CCK-8 solution was added directly to each well (20 µL, i.e. 10% of the culture volume) without removing the medium, and plates were incubated at 37 °C for 0.5h for 293T, DF1 and THP-1 and 2h for PGCs. The absorbance at 450 nm was then measured using a microplate reader. Each condition was tested in 3–5 replicate wells, and background absorbance (media plus CCK-8 in wells without cells) was subtracted. Relative cell viability (%) was calculated by normalizing the absorbance of treated wells to that of control wells (after background correction). For example, viability = 100% × [(A_treated – A_blank) / (A_control – A_blank)]. Growth curves were generated by performing CCK-8 assays at multiple time points (e.g. daily) and plotting the increase in absorbance, which reflects cell proliferation. In addition, cell counts and viability were independently verified using a Trypan Blue exclusion assay on a DeNovix CellDrop automated cell counter. These assays confirmed that PGCs maintained high viability (>90%) in standard culture and that cytotoxic treatments (such as high-dose etoposide) reduced viable cell numbers in a dose-dependent manner.

### Cell Cycle Detection

PGCs were washed once with PBS and collected by centrifugation at 1,500 rpm for 5 min. The cell concentration was adjusted to 1×10⁶/ml, and 1 ml of single-cell suspension was prepared. After centrifugation, the supernatant was discarded, and the cells were fixed with 500 μl of pre-cooled 70% ethanol for 2 hours to overnight at 4°C. Prior to staining, fixed cells were washed with PBS to remove ethanol. The cell pellet was resuspended in 100 μl RNase A solution and incubated at 37°C for 30 min. Then, 400 μl propidium iodide (PI) staining solution (CA1510, Solarbio) was added, followed by incubation in the dark at 4°C for 30 min. Samples were subjected to flow cytometry analysis, and red fluorescence was recorded at an excitation wavelength of 488 nm.

### Data Analysis and Statistics

Unless otherwise noted, all quantitative experiments were performed with n ≥ 3 independent biological replicates. Data are presented as mean ± standard error of the mean (SEM). Statistical analyses were carried out using GraphPad Prism 10. For comparisons between two groups, we used two-tailed Student’s *t*-tests. For multi-group data, one-way or two-way ANOVA was applied followed by appropriate post-hoc tests. *P* < 0.05 was considered statistically significant. All figures report significance levels as *P* < 0.05 (∗), *P* < 0.01 (∗∗), *P* < 0.001 (∗∗∗), and *P* < 0.0001 (∗∗∗∗). Where relevant, outliers were identified using the Grubbs’ test and excluded.

## Results

### Long-term in vitro culture and characterization of PGCs

We successfully established long-term cultures of blood-derived chicken PGCs (PGCs) from 2.5-day embryos, maintaining their germline characteristics in vitro. After ∼3 weeks of culture, contaminating blood cells disappeared and discrete colonies of PGCs emerged in both male (ZZ) and female (ZW) cultures (Figure. 1a). These cells exhibited the expected morphology of avian PGCs – large, refractile cells often in clumps – similar to previous reports of cultured chicken PGCs (Karagenç et al., 1996; Whyte et al., 2015). Growth curve analysis showed that both male and female PGCs entered logarithmic growth by ∼5 days of culture, then reached a plateau by ∼10 days (likely due to the 96-well culture space limitation), with no significant sex-specific difference in proliferation rate (Figure. 1b, *p*-value = 0.14). Established PGC lines could be maintained for >50 days in vitro without spontaneous differentiation.

**Figure 1.**
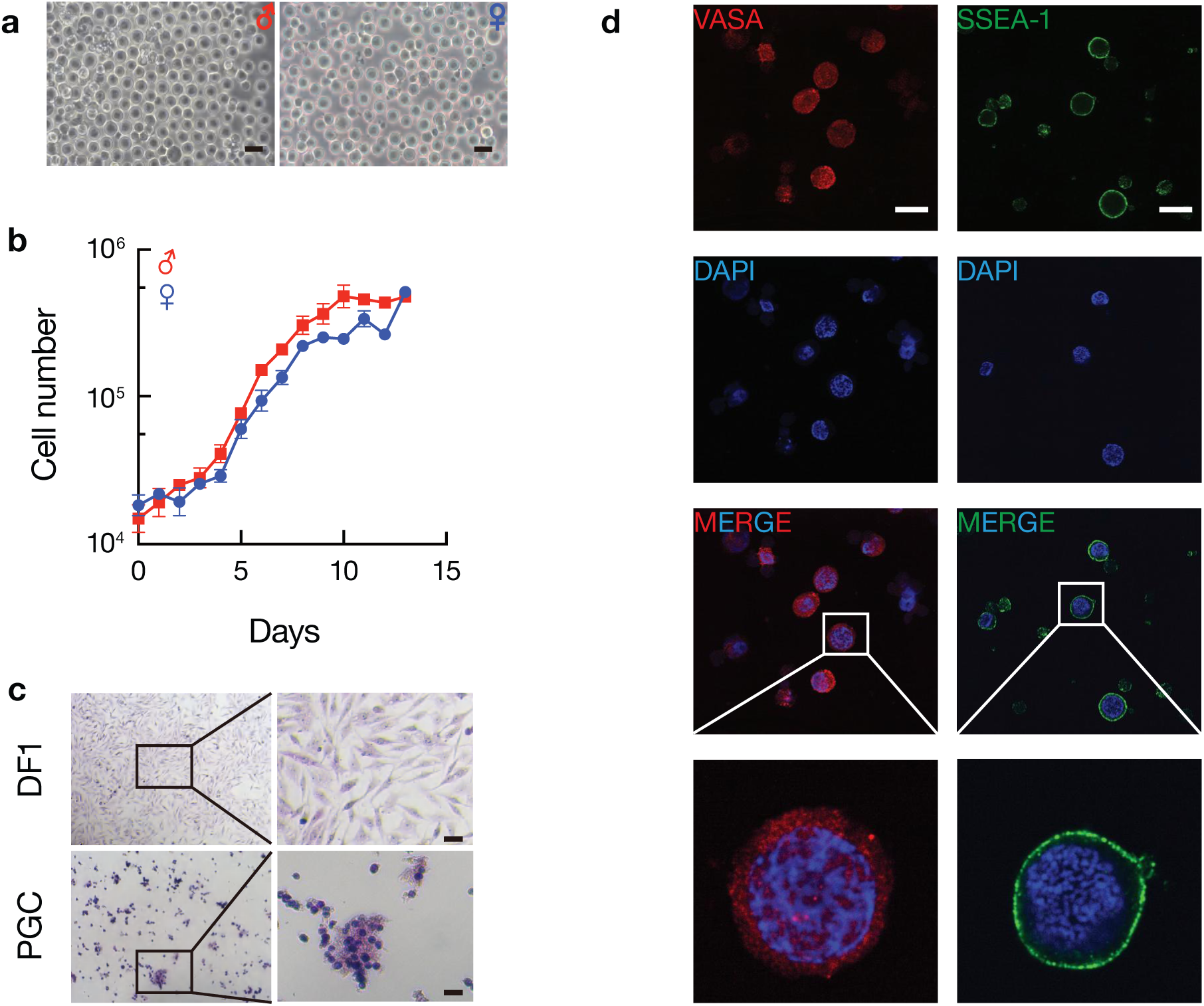
Long-term culture of chicken PGCs in vitro. a) Phase-contrast images of cultured PGCs isolated from embryonic blood, showing typical colony morphology after 3 weeks in vitro. Left: male PGC colony; right: female PGC colony. Scale bar = 20 µm. b) Growth curves of male and female PGC lines. Cell counts (log scale) were measured over 12 days; both sexes enter log-phase growth by Day 5 and plateau by ∼Day 10 due to confluency in 96-well plates. Points represent mean ± SD (n = 3 wells per time point). c) PAS staining of DF1 fibroblasts (upper) versus PGCs (lower). Magenta staining indicates glycogen. PGCs show abundant PAS-positive glycogen granules in the cytoplasm (magenta puncta around the nucleus), whereas DF1 cells show minimal PAS reactivity. Scale bar = 20 µm. d) Immunofluorescence for germ cell markers in PGCs . PGCs (male line shown) strongly express SSEA-1 (green, cell surface) and VASA/DDX4 (red, cytoplasmic), even after long-term culture (50+ days). Nuclei are counterstained with DAPI (blue). Scale bar = 20 µm.

To verify the germ cell identity of our cultured PGCs, we assessed classic PGC markers and features. Periodic Acid–Schiff (PAS) staining revealed abundant magenta-stained cytoplasmic glycogen granules in PGCs, whereas control DF-1 fibroblast cells showed negligible PAS reactivity (Figure. 1c). This is consistent with the known enrichment of glycogen in avian PGC cytoplasm (Meyer, 1960), a feature used continuously to identify PGCs in early chick embryos (Mathan et al., 2023). Immunofluorescence further confirmed that even after ∼50 days in culture, PGCs continued to express the germ cell markers DDX4 (VASA) and SSEA-1, with DDX4 localized to the cytoplasm and SSEA-1 on the cell surface (Figure. 1d). In contrast, DF1 somatic cells were negative for these markers (Supplementary Fig. 1a). The persistence of SSEA-1 and VASA in long-term cultured cells agrees with previous studies showing that cultured chicken PGCs retain germline marker expression over extended periods (Mathan et al., 2023; Yousefi Taemeh et al., 2025). Taken together, these results demonstrate that we established stable male and female PGC lines in vitro that proliferate robustly and maintain key PGC features, including glycogen accumulation and germline-specific protein expression.

### CRISPR/Cas9-mediated EGFP disruption in PGCs and associated toxicity

Having established PGC cultures, we next sought to test gene editing approaches in these cells. We created a stable EGFP reporter PGC line by integrating a CAG promoter–driven EGFP cassette via the Sleeping Beauty transposon system and FACS-sorting EGFP-positive cells (Supplementary Fig. 2A). This PGC-CAG-EGFP line provided a convenient readout for gene knockout efficiency. We synthesize two guide RNAs (gEGFP1 & 2) previously reported to target EGFP (Shalem et al., 2014). PGC-EGFP cells were then co-electroporated with either Cas9 mRNA or Cas9 protein (at various doses) along with the gRNAs, and editing outcomes were analyzed after 5 days.

Strikingly, although the CRISPR/Cas9 treatment effectively introduced mutations at the target site, it also caused substantial toxicity in PGCs. Morphologically, Cas9/gRNA-electroporated PGCs showed evidence of cell death – many cells became fragmented or translucent compared to controls (Figure. 2a). This effect was dose-dependent: higher doses of Cas9 (3 µg vs. 1 µg) led to more pronounced cell loss (Figure. 2a, left panels for Cas9 mRNA; right panels for Cas9 protein). By contrast, unedited control PGCs (carrying the EGFP transgene but not exposed to Cas9) remained healthy and brightly EGFP-positive. Flow cytometry confirmed a marked reduction in EGFP fluorescence in the CRISPR-edited populations (Figure. 2b). The peak EGFP intensity shifted downward in edited cells relative to control, indicating many cells lost or diminished EGFP expression. Quantitative analysis showed that the percentage of EGFP⁺ cells dropped significantly in the Cas9- treated groups (by roughly 50–80% depending on Cas9 dose), accompanied by a sharp decrease in live-cell yield during flow cytometry gating (Figure. 2b, right panel). The loss of EGFP⁺ cells far exceeded the editing efficiency expected from indels alone, suggesting that many PGCs were dying rather than persisting as GFP-negative knockouts. Indeed, cell viability in the highest Cas9 dose group was visibly poor. These observations are in line with reports that excessive CRISPR-induced DNA double-strand breaks (DSBs) can trigger a p53-dependent toxicity in human cultured cells (Mathan et al., 2023), drastically reducing survival of edited cells.

**Figure 2.**
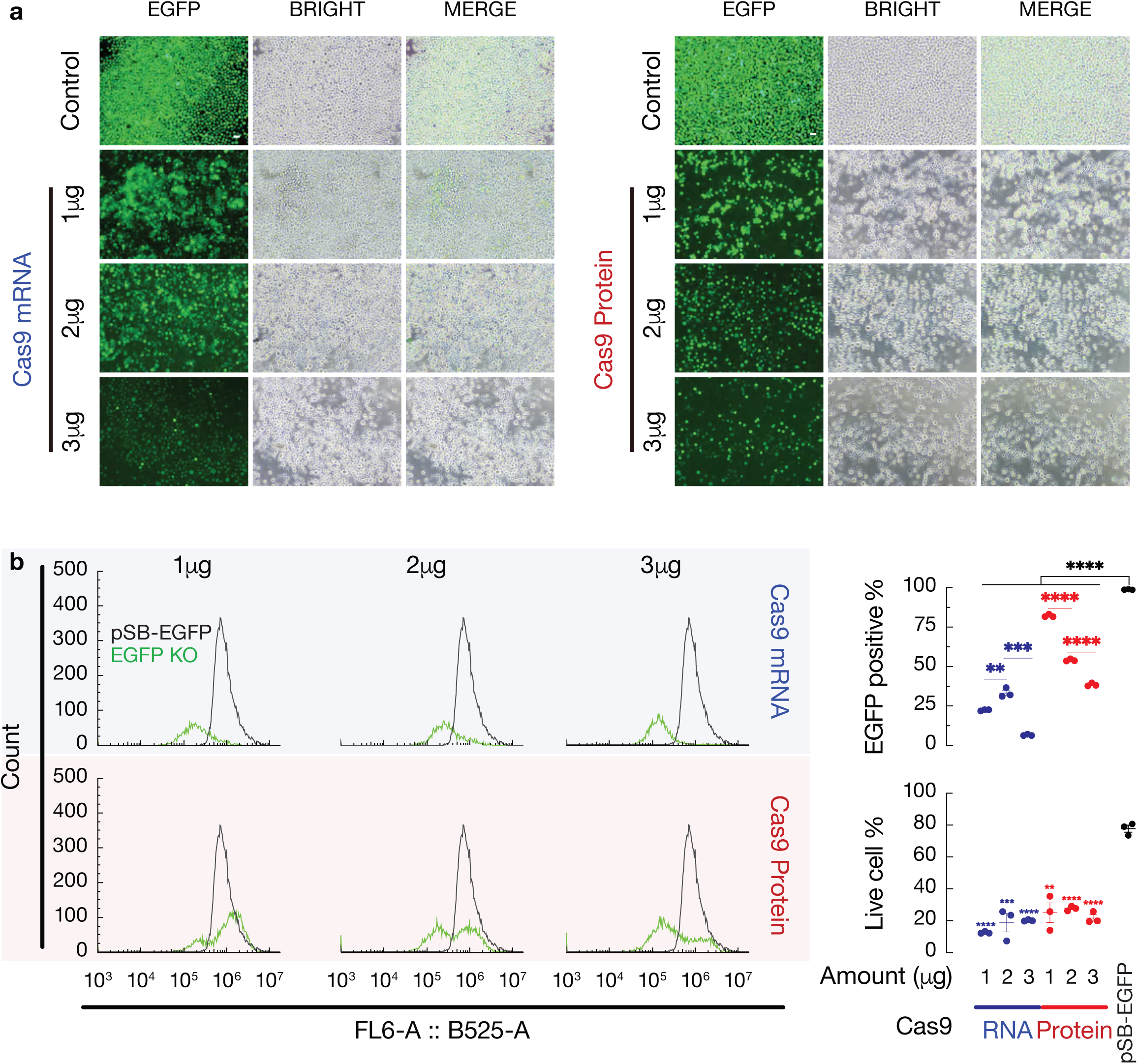
CRISPR/Cas9 editing in PGCs leads to EGFP disruption but reduced cell survival. a) Representative fluorescence microscopy of EGFP⁺ PGCs after co-electroporation with Cas9 and sgRNAs targeting EGFP (gEGFP1+2). **Left:** Cells electroporated with Cas9 mRNA at 1 µg, 2 µg, or 3 µg doses (constant gRNA amount). **Right:** Cells electroporated with Cas9 protein (RNP complex) at equivalent molar amounts (1:1.2 Cas9:sgRNA ratio). Images were taken 5 days post-electroporation. In both RNA and protein conditions, higher Cas9 doses result in loss of EGFP fluorescence and cellular damage (fewer, rounded cells) compared to lower doses. Scale bar = 20 µm. b) Flow cytometry analysis of EGFP fluorescence and cell viability in edited versus control PGCs. **Left:** Histogram overlays of EGFP intensity for control (untreated EGFP⁺ PGCs, gray) vs. CRISPR-edited cells (green). Cas9-edited populations show a shift toward lower fluorescence, indicating EGFP knockout. **Upper right:** Bar graph quantifying the percentage of EGFP⁺ cells in each group (mean ± SD, n = 3). Both Cas9 mRNA and Cas9 protein treatments caused a dose-dependent decrease in the fraction of EGFP-expressing cells compared to control (p<0.01 or p<0.001 by one-way ANOVA). **Lower right:** Viability plot showing the percentage of live cells recovered during flow cytometry. Higher Cas9 doses correlate with reduced live-cell recovery, reflecting CRISPR-induced cytotoxicity in PGCs. Statistical significance was determined by one-way ANOVA (**p<0.01, ***p<0.001, ****p<0.0001).

Molecular analyses verified that the CRISPR/Cas9 treatment successfully introduced indel mutations at the EGFP locus in the surviving cells. Genomic PCR spanning the target site followed by T7 endonuclease I (T7EI) assay yielded cleaved bands in Cas9-edited samples (Supplementary Fig. 2b). The appearance of two smaller fragments (red arrows) indicates heteroduplex formation and cleavage at mismatched DNA, confirming the presence of indels in those cells, whereas the control sample (untreated PGC-EGFP) showed only an intact band (no cutting). Sanger sequencing of the target region, analyzed by the Inference of CRISPR Edit (ICE) deconvolution tool, further demonstrated insertion/deletion mutations around the expected Cas9 cut site (Supplementary Fig. 2c). The ICE analysis revealed diverse indel alleles near the protospacer adjacent motif (PAM) site (dashed red line in Supplementary Fig. 2c) in edited cells, whereas control cells showed the wild-type sequence. These results confirm that our CRISPR/Cas9 approach efficiently introduced mutations in the EGFP reporter gene. However, the concomitant high loss of viable PGCs highlights a major challenge: PGCs appear especially sensitive to the DNA damage caused by CRISPR/Cas9. This aligns with prior findings that while gene-edited PGCs can be obtained (Idoko-Akoh et al., 2018), introducing DSBs may activate DNA damage checkpoints and apoptosis. Our data suggest that standard CRISPR/Cas9 editing in PGCs can exact a steep viability cost, underscoring the need for gentler genome engineering methods in these cells.

### PGCs exhibit heightened sensitivity to DNA damage compared to somatic cells

The pronounced cell death observed following CRISPR editing prompted us to examine the general response of PGCs to DNA damage. We compared the sensitivity of PGCs to that of somatic cells — specifically, HEK 293T (adhesion), DF-1 chicken fibroblasts (adhesion), and THP-1 monocytes (mammalian suspension) — by treating all cell types with etoposide (ETP), a potent topoisomerase II inhibitor that induces DSBs through the stabilization of DNA cleavage complexes (Saleh et al., 2012). Because PGCs were highly vulnerable, we shortened their exposure to 24 hours to avoid excessive loss, while somatic lines were exposed for 48 hours. At low micromolar concentrations of ETP, PGC viability dropped sharply after 24 hours, whereas 293T and DF-1 cells retained more than 50% viability under equivalent or even higher doses over 48 hours; THP-1 cells exhibited intermediate sensitivity (Figure. 3a). These results indicate that PGCs are unusually sensitive to chemically induced DSBs, likely reflecting a heightened germline strategy to eliminate damaged cells rather than risk genomic integrity.

**Figure 3.**
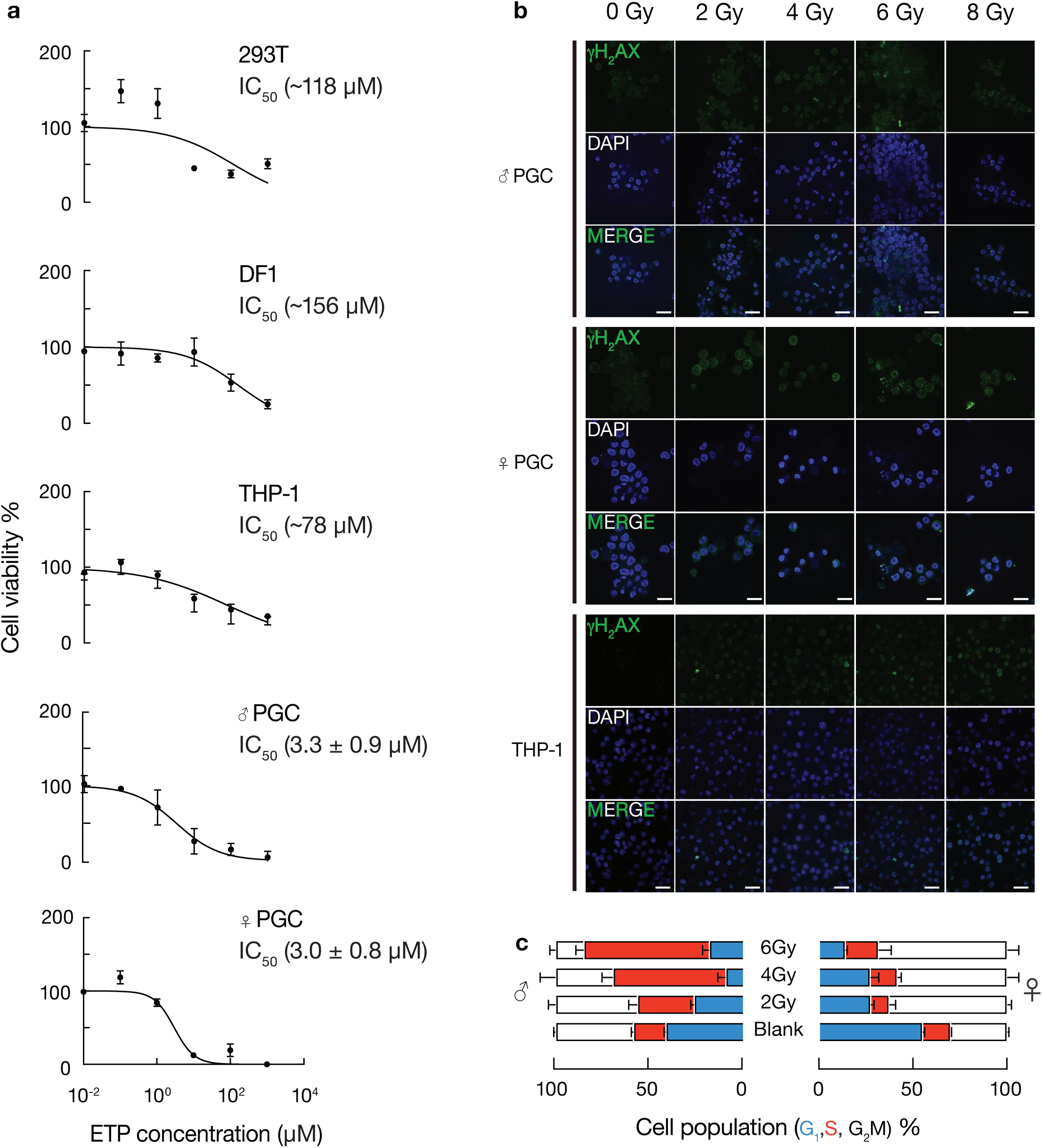
PGCs are more sensitive to DNA double-strand breaks than somatic cells. a) Cell viability after etoposide (ETP) treatment. Dose–response curves (log_10_ [ETP] on x-axis) are shown for 293T and DF1 (48 h treatment), for THP-1 cells and PGCs (24 h treatment). PGC viability drops sharply even at low ETP doses, whereas 293T, DF1, and THP-1 cells are more tolerant to ETP (viability data are mean ± SD of triplicates). b) DNA damage marker immunofluorescence after X-ray irradiation. Shown are representative images of male PGCs (upper), female PGCs (middle), and THP-1 cells (lower) several minutes after exposure to 0, 2, 4, 6, or 8 Gy and recover for 48 h. Green: γ-H2AX foci indicating DSBs; Blue: DAPI nuclear stain. PGCs exhibit γ-H2AX foci at lower doses (2– 4 Gy) than THP-1 cells, and extensive pan-nuclear γ-H2AX at 8 Gy, suggesting less DNA damage tolerance or repair. Scale bar = 50 µm. c) Cell-cycle distribution of male vs. female PGCs after DNA damage. PGCs were irradiated (2 Gy, 4 Gy and 6 Gy), cultured 48 h, and analyzed by flow cytometry for cell-cycle phase (propidium iodide staining). Stacked bar charts show the percentage of cells in G_0_G_1_, S, and G_2_/M phases for untreated vs. irradiated cells. After damage, female PGCs predominantly arrest in G_2_/M (increased G_2_ fraction), whereas male PGCs accumulate in S-phase.

We next exposed cells to ionizing radiation and examined DNA damage and survival. Immunofluorescence staining for phosphorylated H2AX (γ-H2AX), a well-established marker of DSBs (Kinner et al., 2008), was performed on male and female PGCs and, for comparison, on THP-1 cells after 0, 2, 4, 6, or 8 Gy of X-ray irradiation. As shown in Figure 3b, both male and female PGCs showed dose-dependent formation of nuclear γ- H2AX foci (green) after radiation. Notably, at moderate doses (2–4 Gy), PGCs already exhibited numerous γ-H2AX foci, whereas THP-1 cells required higher doses (6–8 Gy) to show comparable levels of foci. This suggests PGCs incur or retain more DSBs per unit of irradiation, or activate the damage signal more readily. By 8 Gy, extensive DNA damage was evident in all cell types, but many PGC nuclei appeared intensely stained, often with pan-nuclear γ-H2AX, indicative of severe or unrepaired damage. Consistent with the immunofluorescence, cell viability assays 48 h post-irradiation showed that PGCs were markedly radiosensitive (Table 1). Increasing radiation doses caused a steep decline in PGC survival, significantly greater than the viability loss observed in 293T or DF1 fibroblasts over the same dose range (THP-1 viability also declined, but these cells are somewhat more radiation-resistant in our assays). For example, at 6 Gy, the majority of PGCs were non-viable, whereas a substantial fraction of the somatic cells were still alive. This heightened sensitivity of PGCs to DSBs is in line with their germline role – to preserve genomic integrity, germ cells may activate checkpoints or apoptosis more readily upon DNA damage.

**Table 1.** Cell viability after X-ray treatments.

| Treatments | 293T | DF1 | THP-1 | ♂ bPGC | ♀ bPGC |
| --- | --- | --- | --- | --- | --- |
| Blank | 100 | 100 | 100 | 100 | 100 |
| 2Gy | 114±28 | 101±7 | 103±4 | 75±9*** | 63±7 |
| 4Gy | 122±14 | 91±7 | 116±17 | 79±2** | 64±13 |
| 6Gy | 120±16 | 107±35 | 109±35 | 63±8**** | 55±29* |
| 8Gy | 115±23 | 75±5 | 79±18 | 56±4**** | 68±15 |

Interestingly, we observed a sex-specific difference in cell-cycle arrest in PGCs following DNA damage. We performed cell-cycle analysis on male and female PGC cultures 48 h after irradiation. In the absence of damage, both male and female PGCs had cell-cycle profiles typical of proliferating cells (substantial S-phase populations and G_2_/M fractions).

After X-ray exposure, however, male and female PGCs accumulated at different phases (Figure. 3c). Female PGCs showed a pronounced G_2_/M arrest – the majority of cells shifted into the G_2_/M phase, with a corresponding reduction in S-phase population. In contrast, male PGCs exhibited a significant accumulation in S-phase and a comparatively smaller increase in G_2_/M. In other words, DNA damage caused most female PGCs to halt before mitosis (perhaps via an ATM/CHK2-mediated G_2_ checkpoint), whereas male PGCs more often slowed or stalled during DNA synthesis (S-phase, possibly via ATR/CHK1 activation). This divergence resulted in a statistically significant difference in cell-cycle distribution between the sexes upon damage (p<0.01 for both S and G_2_/M fractions). The basis for this sex difference is unclear, but it may reflect underlying differences in cell-cycle regulation or DNA repair pathways between ZZ and ZW germ cells. Regardless, both sexes clearly activate checkpoints in response to DSBs, further highlighting that PGCs do not readily tolerate genomic damage. Overall, our findings demonstrate that chicken PGCs are highly sensitive to DNA double-strand breaks, responding with robust DNA damage signaling (γ-H2AX) and cell-cycle arrest or apoptosis at levels of damage that somatic cells can partially withstand.

### CRISPRi versus shRNA for gene silencing in chicken cells

Finally, we evaluated alternative, potentially less genotoxic methods for gene perturbation in PGCs and chicken cells. CRISPR interference (CRISPRi), which uses a catalytically dead Cas9 (dCas9) fused to a repressor domain to epigenetically silence target genes (Gilbert et al., 2013), has proven effective in mammalian cells. We tested whether CRISPRi could similarly repress gene expression in chicken cells without inducing DNA breaks. We co-transfected cells with two plasmids: (1) a PGK-CRISPRi-EGFP construct expressing the dCas9-KRAB fusion protein (along with an EGFP marker) under a constitutive promoter, and (2) a gCAG-mCherry construct expressing an sgRNA (targeting the CAG promoter sequence) from the chicken U6.3 promoter and a CAG-driven mCherry reporter (Figure. 4a). The design was to target the same CAG promoter driving mCherry in our reporter cells, thus CRISPRi should knock down mCherry transcription. As a positive control for CRISPRi function, we performed parallel experiments in human 293T cells carrying a similar CAG-mCherry reporter.

**Figure 4.**
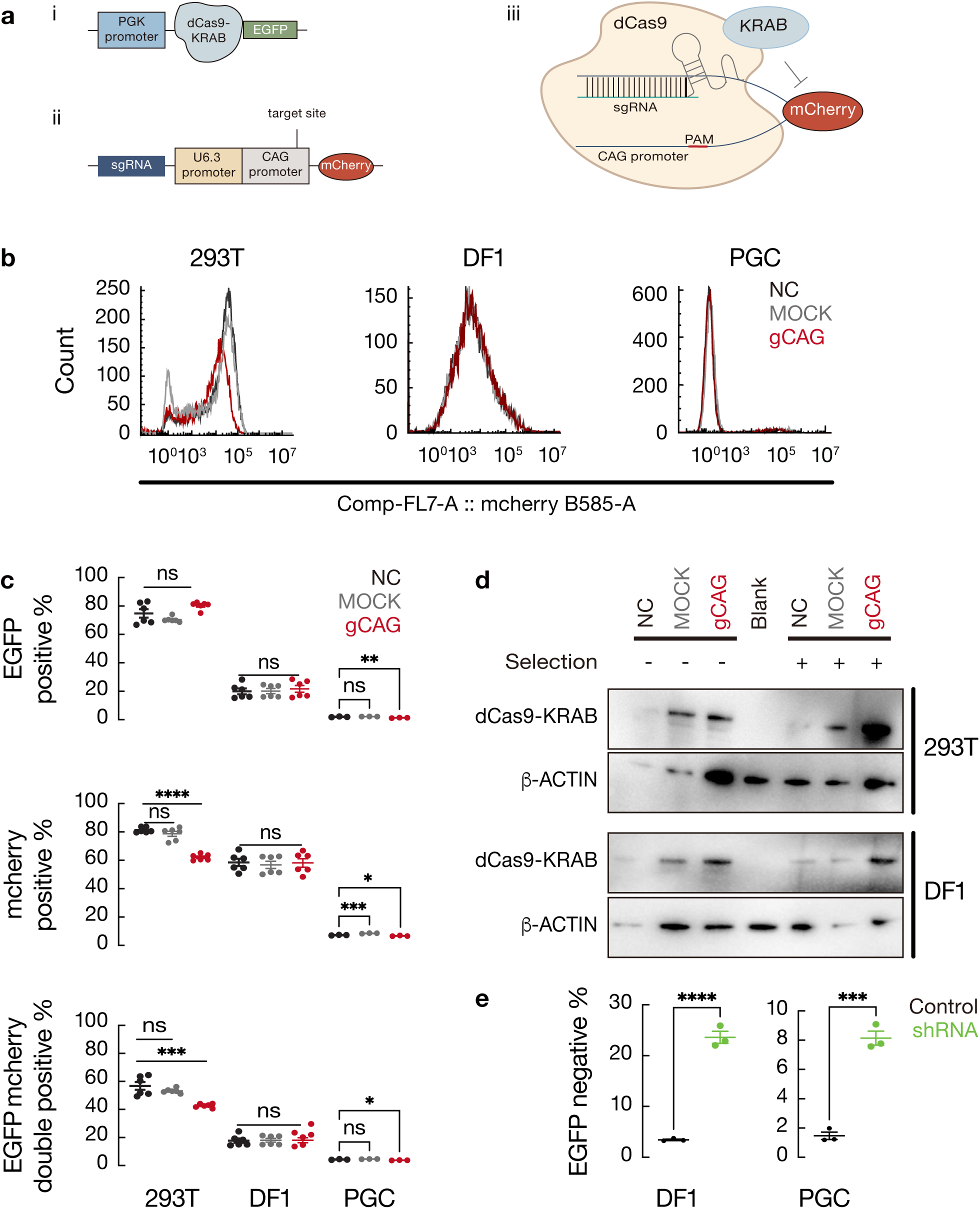
CRISPRi is less effective in chicken cells than in human cells, whereas shRNA knockdown is effective in both. a) Schematic of the CRISPR interference (CRISPRi) system used. (i) The PGK-CRISPRi-EGFP plasmid expresses dCas9-KRAB (a fusion of catalytically inactive Cas9 and the KRAB repressor domain) and an EGFP marker under a constitutive PGK promoter. (ii) The gCAG-mCherry plasmid carries a U6.3 promoter-driven sgRNA targeting the CAG promoter and a CAG-driven mCherry reporter. (iii) Co-transfection strategy: dCas9-KRAB (from plasmid i) is expressed in the cell, and the sgRNA (from plasmid ii) guides it to the CAG promoter present in the mCherry cassette, ideally silencing the transcription of mCherry. b) Flow cytometry histograms of mCherry fluorescence in cells transfected with the gCAG-mCherry vector. Black line: cells lacking sgRNA (empty vector control); gray filled: mock transfected cells; red line: cells with the CAG-targeting sgRNA. **Left:** 293T human cells expressing dCas9-KRAB. The red histogram is slightly shifted left (lower mCherry intensity) compared to black, indicating CRISPRi-mediated repression of mCherry expression. **Middle:** DF1 chicken cells expressing dCas9-KRAB. The mCherry fluorescence profiles overlapped across all conditions. There was no discernible decrease in the presence of the targeting sgRNA (red vs. black), suggesting dCas9-KRAB did not effectively silence the CAG promoter in DF1 cells; similar results were observed in PGCs (**Right**). c) Summary of CRISPRi reporter knockdown efficacy in 293T vs. chicken cells. **Upper:** Percentage of EGFP⁺ cells (transfection efficiency) in each condition. High and similar EGFP⁺ percentages in both cell types confirm effective delivery of the CRISPRi plasmid; small differences between conditions reflect only transfection variability. **Middle:** Percentage of mCherry⁺ cells in each condition (no sgRNA, mock, +gCAG sgRNA) for human vs. chicken cells. In 293T, introduction of the CAG-targeting sgRNA reduces the mCherry⁺ fraction relative to controls, whereas in DF1 the mCherry⁺ percentage remains unchanged. In PGC, there is a small but statistically significant decrease in mcherry expression. **Lower:** Percentage of double-positive (EGFP⁺ mCherry⁺) cells. In 293T, the CAG-targeting sgRNA significantly lowers the double-positive fraction (indicating many mCherry-transfected cells lost mcherry expression), while in DF1 there is no significant change. PGC shows a weak reduction in double-positive fraction. Data are mean ± SD (n = 3); statistical comparisons by one-way ANOVA, p<0.01. d) Western blot analysis of dCas9-KRAB protein expression in 293T and DF1 CRISPRi-expressing cells, before (–) and after (+) puromycin selection. A specific band for dCas9-KRAB (∼205 kDa) is detected in both cell types after selection (plus lanes). β-ACTIN is shown as a loading control. The blot confirms that lack of CRISPRi efficacy in DF1 was not due to an absence of dCas9-KRAB expression. e) shRNA-mediated EGFP knockdown in chicken and human cells. DF1-CAG-EGFP and 293T-CAG-EGFP reporter cells were transfected with a plasmid encoding an shRNA against EGFP. The percentage of EGFP⁺ cells was measured 4 days post-transfection by flow cytometry and normalized to no-treated EGFP⁺ cells. Both DF1 and PGCs cells showed a significant reduction in EGFP⁺ cells following EGFP shRNA introduction (∼20– 30% of cells lost EGFP in DF1, ∼8% of cells lost EGFP in PGCs). Data represent mean ± SD (n = 3); **p**<0.01 by two-tailed t-test comparing EGFP shRNA vs. control. (In panels c, e: \***p**<0.01, \*\***p**<0.001, \*\*\***p**<0.0001).

Despite robust dCas9-KRAB expression, CRISPRi did not achieve appreciable gene knockdown in chicken cells. In transient transfections, the gCAG mCherry vector was introduced into PGCs, DF1, and 293T cells that express dCas9-KRAB. mCherry fluorescence served as an indicator of sgRNA vector uptake. In 293T cells, we observed that introducing the CAG-targeting sgRNA caused a subtle shift in mCherry fluorescence intensity (Figure. 4b), consistent with dCas9-KRAB binding to and partially repressing the CAG promoter driving mCherry on the same plasmid. In chicken cells (DF1 and PGCs), however, the mCherry fluorescence distribution was virtually unchanged whether a targeting sgRNA was present or not (Figure. 4b), suggesting dCas9-KRAB was not effectively silencing the CAG promoter in these cells. Flow cytometry quantification across replicate experiments showed that in 293T cells co-transfected with CRISPRi (dCas9-KRAB) and gCAG, the percentage of mCherry-positive cells dropped (and the mCherry mean fluorescence intensity was lower) compared to controls, whereas in DF1, the mCherry expression remained high and nearly identical to controls. PGC showed a slight but significant decrease in mcherry expression (Figure. 4c). Specifically, the fraction of EGFP⁺/mCherry⁺ double-positive cells was significantly reduced in 293T (indicating successful mCherry repression in those cells that received the sgRNA), but no significant reduction was seen in DF1 while in PGCs, there is a weak reduction in double-positive cells (Figure. 4c, right panel). These results suggest that the CRISPRi system, while functional in a human context, feeble to repress the target gene in chicken cells under our conditions.

We pursued this further by establishing stable cell pools expressing dCas9-KRAB and the gCAG sgRNA (via antibiotic selection of co-transfected cells). Even after selection – which ensured that essentially all cells carried dCas9-KRAB and the sgRNA plasmid – the CAG promoter was still strongly silenced in the human cells, not the chicken cells. In 293T pools, the mCherry fluorescence profile shifted markedly to lower intensity when the CAG-targeting sgRNA was present, reflecting CRISPRi-mediated repression of the CAG promoter driving mCherry (Supplementary Fig. 3a, left). In contrast, DF1 pools with the same sgRNA showed no shift in mCherry levels relative to control, indicating a lack of CAG promoter repression. Quantification revealed that a large proportion of 293T cells (>50%) exhibited a >10-fold drop in mCherry fluorescence with the targeting sgRNA, whereas virtually no DF1 cells did (Supplementary Fig. 3a, right). Importantly, Western blot confirmed that dCas9-KRAB protein was expressed in both the selected DF1 and 293T cells at comparable levels (Figure. 4d). Thus, the inefficient of CRISPRi in chicken cells was not due to a lack of repressor protein. Instead, it likely reflects an inherent limitation of the CRISPRi mechanism in the avian chromatin context – for example, the chicken KRAB–KAP1 repression pathway or sgRNA expression from the U6.3 promoter may not be as effective. Our findings concur with a recent study that reported only modest (∼50– 60%) gene downregulation using CRISPRi in chicken DF1 cells (Chapman et al., 2023), whereas CRISPRi in mammals often achieves >90% knockdown (Gilbert et al., 2013). In sum, dCas9-KRAB-mediated transcriptional repression appears to be markedly less efficient in chicken cells than in human cells.

Given the shortcomings of CRISPRi in our hands, we turned to RNA interference as an alternative knockdown strategy. Short hairpin RNA (shRNA) plasmids targeting EGFP were transfected into the CAG-EGFP reporter cells, and gene silencing was evaluated by flow cytometry. In stark contrast to CRISPRi, the shRNA approach was highly effective in chicken cells. A few days after transfection of an anti-EGFP shRNA, the percentage of EGFP^-^ cells increased dramatically in DF1 cultures, from nearly 3% to 20–30% (Figure. 4e and Supplementary Fig. 3b). This degree of silencing was statistically significant (p<0.0001 vs. control shRNA) and corresponded to a roughly 20 % reduction in EGFP⁺ cells. Such efficiency is comparable to knockdown levels reported for shRNAs in chicken fibroblasts by others (Guru Vishnu et al., 2019). Even in PGC, nearly 8% cells lost high intensity EGFP fluorescence. Thus, conventional RNAi remains a reliable method for gene suppression in chicken PGCs or fibroblast cells. In summary, our results indicate that while CRISPR/Cas9 genome editing can induce mutations in PGCs, it does so at the cost of substantial cell damage, and CRISPR-based transcriptional repression (CRISPRi) that works well in mammals is inefficient in the avian system. By comparison, shRNA-mediated knockdown caused no overt toxicity and achieved robust gene silencing in chicken cells. These findings underscore the need to optimize or choose alternative genetic manipulation tools for germline cells in birds, with shRNA emerging as a preferable approach for functional studies in chicken PGCs.

## Discussion

### Germline DNA Damage Sensitivity and Sex-Specific Responses

Germ cells are uniquely sensitive to DNA damage, a trait conserved across species and thought to safeguard genomic integrity for future generations. In both mice and humans, primordial germ cells (PGCs) are hypersensitive to endogenous and exogenous DNA damaging agents(Bloom and Schimenti, 2020). This extreme sensitivity of the germline to genotoxic stress is well documented in many organisms, from insects to mammals(OAKBERG, 1955; Arnon et al., 2001; Meistrich, 2013; Gunes et al., 2015; Lu and Yamashita, 2017). Evolutionarily, this likely functions as a quality control mechanism to prevent transmission of mutations: damaged germ cells are eliminated rather than repaired imperfectly. Indeed, unlike most somatic cells which attempt cell cycle arrest and repair after DNA damage, germ cells (and pluripotent stem cells) often favor apoptosis over error-prone repair, thereby avoiding the risk of propagating deleterious mutations(Bloom et al., 2019).

Notably, DNA damage can severely impact fertility, underscoring germ cells’ vulnerability. In female mammals, the oocyte pool is finite and highly susceptible to depletion by radiation or chemotherapy(Arnon et al., 2001; Meistrich, 2013; Gunes et al., 2015; Lu and Yamashita, 2017). This is exemplified by the DNA damage response in oocytes, where even a few double-strand breaks can trigger robust activation of the p53 family protein TAp63 and consequent apoptosis, rapidly culling damaged oocytes to preserve species fitness(Lu and Yamashita, 2017). The high sensitivity of female germ cells is partly explained by their limited reserve – any loss is irrecoverable – whereas males continually produce sperm. Nonetheless, male germ cells also show pronounced DNA damage sensitivity, a phenomenon not fully explained by numbers alone. One hypothesis is that additional mechanisms (such as intercellular germ cell connectivity in cysts) amplify damage signals to ensure even subtle DNA lesions in spermatogonia lead to coordinated cell death(Lu and Yamashita, 2017). This concerted response, observed in Drosophila and presumably in mammals, helps maintain the genetic integrity of the gamete pool by sacrificing compromised cells for the sake of offspring genome quality. In summary, germline cells across species have evolved a “zero-tolerance” strategy for DNA damage – an extreme vigilance that, while protective for the next generation, carries the trade-off of high fertility risk under genotoxic stress (as seen in cancer therapies and environmental exposures).

Importantly, there are distinct sex-dependent nuances in these responses. Female germ cells (oocytes) undergo stringent quality control; for example, TAp63-mediated checkpoints purge DNA-damaged oocytes to prevent mutated embryos(Lu and Yamashita, 2017). This makes female fertility especially vulnerable, but it is an essential safeguard against transmitting genetic errors. Male germ cells, despite their renewable nature, also exhibit damage-induced attrition (e.g. spermatogonial apoptosis) albeit via potentially different molecular pathways (p53, intrinsic checkpoints, etc.). The male germline’s sensitivity is more puzzling given continuous sperm production, but it likely reflects conserved surveillance mechanisms that favor genomic integrity over cell survival(Lu and Yamashita, 2017). Thus, both sexes’ germ cells are hyper-vigilant, but the consequences differ: in females, it manifests as irreversible loss of primordial follicles with age or insult, whereas in males, it may lead to transient infertility or reduced sperm quality. These differences underscore the influence of germ cell biology – e.g. finite vs. self-renewing germ cell pools – on how DNA damage responses are tuned. Understanding these sex-specific responses has practical implications, such as developing fertility-preservation strategies (e.g. protecting oocytes during chemotherapy) and informing germline engineering approaches (which must contend with the germ cells’ propensity to undergo apoptosis if genome integrity is threatened).

### Overcoming Challenges in Chicken PGC Genome Editing

Compared to mammals, the progress in avian genome editing has been relatively slow(Lee et al., 2020a), in part due to technical hurdles and the sensitivity of avian germ cells. Cultured chicken PGCs provide a valuable route to genetically modify the germline, but efficient transfection and editing have historically been difficult. In our study, we addressed this by optimizing a delivery method for CRISPR/Cas9 in chicken PGCs. Specifically, we developed an electroporation-based approach to introduce Cas9 mRNA or ribonucleoprotein, which proved highly effective and less toxic than conventional lipid-based transfection. Consistent with a recent optimization report, we found that electroporation can achieve high transfection efficiencies (on the order of 50–70% of cells expressing the transgene) while maintaining normal PGC growth and pluripotency marker expression(Zhao et al., 2025). Lipofection of plasmid DNA in PGCs, by contrast, yielded lower uptake and higher cell stress in our hands, whereas electroporation (particularly using a nucleofection device and carefully tuned pulse parameters) resulted in minimal apoptosis and robust gene integration or editing. By delivering Cas9 as mRNA or protein rather than via persistent plasmid expression, we likely avoided prolonged nuclease activity, thereby reducing DNA damage responses in the PGCs.

An additional challenge in genome editing of chicken PGCs is the potential toxicity of Cas9 itself. High, constitutive expression of Cas9 has been reported to be detrimental in various systems – for example, attempts to create Cas9-transgenic chickens showed that embryos homozygous for a Cas9 insertion often failed to hatch, likely due to toxicity from excessive Cas9 activity(Duda, 2021). Even in mammalian cells, a single DNA double-strand break induced by Cas9 can trigger p53-dependent toxicity in pluripotent stem cells(Kim et al., 2014; Idoko-Akoh et al., 2018). Thus, the sensitivity of PGCs to DNA breaks means that standard CRISPR/Cas9 editing (which creates targeted DSBs) can lead to cell cycle arrest or apoptosis, undermining editing efficiency. Our electroporation strategy mitigates some of this by using transient Cas9 RNP delivery, thereby limiting the duration of Cas9 activity and off-target DSB formation. We also foresee that using high-fidelity or “nickase” variants of Cas9 could further reduce unintended damage.

Looking ahead, strategies that avoid making double-strand breaks entirely may be especially advantageous for editing the germline of chickens and other sensitive cell types. Base editors and prime editors represent two such innovations that perform precise nucleotide conversions with either single-strand nicks or engineered reverse transcription, rather than blunt DSBs. These newer genome editing tools could circumvent the pronounced DNA damage response of PGCs. Indeed, recent studies have shown promise using base editing in chicken PGCs – for example, introducing precise point mutations in genes like *TF* or *MSTN* without detectable cytotoxicity, and successfully generating germline-transmitting chimeric chickens from those base-edited PGCs(Lee et al., 2020b). Prime editing has likewise been demonstrated in cultured chicken cells, including PGCs, enabling precise insertions/deletions or substitutions without relying on host HDR pathways(Atsuta et al., 2022; Kim et al., 2023). While still emerging technologies, base and prime editing could mitigate the low survival rates associated with editing by sidestepping the need for error-prone end joining and thereby avoiding activation of apoptosis pathways. We suggest that future work in avian genome engineering invest in adapting such DSB-free editors to chicken cells. By reducing genotoxic stress on PGCs, these approaches may achieve higher editing efficiencies and cell viabilities, ultimately accelerating the production of transgenic or gene-edited chickens. In summary, our experience underscores that both the delivery method and the choice of genome editing modality are critical for success in chicken PGCs: electroporation of transient Cas9 or the use of gentler editing enzymes can overcome past limitations and improve the feasibility of precise germline modifications in avian species.

### CRISPR Interference Limitations in Avian Cells

Besides gene knockout, we explored CRISPR interference (CRISPRi) as a means of gene knockdown in chicken PGCs. CRISPRi uses a catalytically inactive dCas9 fused to repressor domains to epigenetically silence target genes. In theory, this approach can achieve tunable suppression of gene expression without permanent genomic alterations or triggering DNA damage. However, our results with CRISPRi in chicken germ cells were unexpectedly underwhelming – we observed little reduction in target gene expression, similar to prior reports in other systems that showed partial knockdown (Chapman et al., 2023), not as strong as in mammalian cells. There are several possible explanations for this discrepancy. One factor is the assay sensitivity and the expression level of the target genes.

If a target gene is very highly expressed or has multiple transcription start sites, a modest repression by CRISPRi might not register as statistically significant downregulation in our assay. For instance, the study applying CRISPRi in chicken DF-1 fibroblasts noted that certain genes (like high mobility group A1 [*HMGA1*] and subfamily b, member 1 [*SMARCB1*]) could not be appreciably knocked down by dCas9-KRAB, potentially because of their high basal expression or regulatory redundancy (Chapman et al., 2023). In our hands, it is possible that the genes tested have similar characteristics, or that the timing of analysis missed the window of maximal repression. We also recognize that CRISPRi efficiency heavily depends on guide RNA positioning; guides must target proximal to the transcription start site (within a window of approximately –50 to +300 bp) for optimal interference. If our gRNA designs were suboptimal or if the chicken gene annotations for transcription start sites were incomplete, we may have inadvertently used less effective guide locations. Differences in promoter context or chromatin accessibility in PGCs (which have a unique epigenetic state during development) could further influence CRISPRi efficacy.

Another major consideration is the design of the CRISPRi construct – specifically, the choice of repressor domains – and its compatibility with avian transcriptional machinery. CRISPRi systems have evolved in mammalian cell studies, where the dCas9 is often fused to human repressor domains such as KRAB (from KOX1) and sometimes additional domains like MeCP2 or others to enhance potency(Duke et al., 2020; Usluer et al., 2023). We used a dCas9-KRAB-based repressor (with a mammalian KRAB domain) in our experiments. Interestingly, a recent comparative study in chicken cells found that a dCas9- KRAB fusion was effective at repressing certain chicken genes (e.g. achieving significant knockdown of *IRF7* when targeting its promoter)(Chapman et al., 2023). However, the same study noted that a dCas9-KRAB-MeCP2 fusion – which in human cells is considered a more potent second-generation CRISPRi – failed to enhance repression in chicken cells. The authors attributed this to the divergence in transcriptional co-repressors between species: the MeCP2 domain (derived from rat in that construct) interacts with mammalian co-repressors like DNMT1 and Sin3A/HDAC complex, but the chicken MeCP2 protein shares only ∼42% identity with the rodent version, likely limiting cross-species functionality. In essence, the epigenetic silencing toolkit of mammals may not fully operate in birds. It is plausible that the human KRAB domain we used also has suboptimal affinity for chicken KAP1 or other co-factors, reducing the efficacy of dCas9-mediated heterochromatin formation. Therefore, the weak of CRISPRi effect in our chicken PGCs could stem from using repressor modules that are not “tuned” to the avian epigenome. Differences in codon usage or expression levels of the dCas9 fusion could further contribute; if the dCas9-KRAB protein was not expressed at high enough levels in PGCs or not properly localized to the nucleus, its impact would be minimal.

Our findings highlight that CRISPRi technology, as currently configured, is not readily transferable across species without modification. To achieve reliable CRISPR-based gene repression in chickens, new tools may need to be developed. For example, one could screen for chicken-derived repressor domains or adapt orthologous silencing proteins from avian sources to fuse to dCas9, ensuring they recruit the correct chicken co-repressor complexes. Engineering the dCas9 protein or the guide RNA scaffolds to better engage avian histone modifiers is another potential route. Until such chicken-optimized CRISPRi systems are available, researchers may find that traditional knockdown methods (e.g. RNA interference) are more dependable for suppressing gene function in avian cells. This was borne out in our study: while CRISPRi had little effect, shRNA-mediated knockdown in PGCs was effective, indicating that the RNAi pathway in chickens can be harnessed for gene silencing. In the context of functional genomics, one should therefore approach CRISPRi in non-mammalian species with caution – validation in the target species is essential, and alternative strategies should be considered when CRISPR-based modulation fails to yield a phenotype.

In conclusion, this study successfully establishes a robust long-term culture system for blood-derived primordial germ cells (PGCs) in chickens, which consistently maintain key germline markers and stable proliferation capacity over extended periods. Notably, conventional CRISPR/Cas9-mediated editing in these cells induces severe DNA damage-dependent toxicity, leading to substantial cell death despite successful mutagenesis. Further comparative analysis reveals that PGCs exhibit significantly heightened sensitivity to DNA double-strand breaks compared to somatic cells, accompanied by sexually dimorphic cell-cycle checkpoint responses. Given these constraints, we evaluated CRISPR interference (CRISPRi) as a potentially gentler alternative, but found it inefficient for transcriptional repression in avian cells, likely due to species-specific limitations in epigenetic silencing machinery. In contrast, shRNA-mediated knockdown achieved efficient and nontoxic gene silencing without triggering DNA damage. Collectively, these findings highlight the unique vulnerability of avian germ cells to genomic perturbation and underscore the necessity of adopting non-nuclease-based approaches, such as RNAi, for functional genetic studies in chicken PGCs.

## Data availability

The data underlying this article are available in the article and in its online supplementary material.

## Acknowledgments

We extend our sincere gratitude to the Public Platform of the International Health Medicine Institute of Zhejiang University for their invaluable assistance with the experimental equipment. We are deeply thankful to Professor Jeffery Barrow, Associate Professor Li Yang, Associate Researcher Deivendran, Associate Researcher Zhao Jingrong, Li Guangnan, and all the members of the Li laboratory for their insightful discussions and contributions throughout the course of this research.

## Supplementary Materials

Supplementary Figure 1-3

Table S1

